# Interpretable Trajectory Inference with Single Cell Linear Adaptive Negative-binomial Expression (scLANE) Testing

**DOI:** 10.1101/2023.12.19.572477

**Authors:** Jack R. Leary, Xiaoru Dong, Rhonda Bacher

**Affiliations:** Department of Biostatistics, University of Florida

**Keywords:** genomics, trajectory analysis, scRNA-seq

## Abstract

The rapid proliferation of trajectory inference methods for single-cell RNA-seq data has allowed researchers to investigate complex biological processes by examining underlying gene expression dynamics. After estimating a latent cell ordering, statistical models are used to determine which genes exhibit changes in expression that are significantly associated with progression through the biological trajectory. While a few techniques for performing trajectory differential expression exist, most rely on the flexibility of generalized additive models in order to account for the inherent nonlinearity of changes in gene expression. As such, the results can be difficult to interpret, and biological conclusions often rest on subjective visual inspections of the most dynamic genes. To address this challenge, we propose scLANE testing, which is built around an interpretable generalized linear model and handles nonlinearity with basis splines chosen empirically for each gene. In addition, extensions to estimating equations and mixed models allow for reliable trajectory testing under complex experimental designs. After validating the accuracy of scLANE under several different simulation scenarios, we apply it to a set of diverse biological datasets and display its ability to provide novel biological information when used downstream of both pseudotime and RNA velocity estimation methods. scLANE is freely available as an R package and is also accessible via a web server leveraging high performance computing resources at https://sclane.rc.ufl.edu/.

## 1 Introduction

Single-cell RNA sequencing (scRNA-seq) has advanced our ability to obtain high-resolution insights into dynamic biological processes including cellular differentiation and disease progression [1]. By capturing sufficient cellular heterogeneity representing progression through a biological process of interest, a cellular trajectory can be computationally inferred based on variations among cells transcriptomic profiles. Such techniques, referred to as pseudotime or trajectory inference methods, estimate a latent cellular ordering that approximates a temporal progression through an underlying process. Trajectories need not be linear and, more generally, processes are constructed as graph-like structures divided into various lineages indicating distinct subprocesses. Trajectory inference methods have successfully identified major transitions in disease [2], cell cycle [3], and development [4]. The trajectory itself, though, is not the final goal. Once a trajectory is constructed, researchers seek to understand which genes are dynamic, and then characterize how those genes’ expression changes along pseudotime.

Trajectory differential expression (DE) analysis aims to identify significantly dynamic genes, and ideally, enable comparisons of gene dynamics between lineages and experimental conditions. Biologically relevant details such as locations along pseudotime where transcriptional switches occur as well as interpretable slope estimates would allow for quantitatively comparing dynamics across genes or lineages. However, in practice, these comparisons are still largely performed visually, or limited to *p*-values providing no interpretable quantitative outputs describing a gene’s expression pattern. Given that scRNA-seq analyses are often exploratory in nature, researchers stand to benefit from statistics that offer a clearer biological context. In addition, with experiments growing more complex, containing multiple subjects and often timepoints [1], flexible methodologies are needed to accommodate a variety of experimental designs and biological questions.

Several single-cell specific methods have been proposed for trajectory DE testing [5–10]. However, all existing approaches lack either interpretability or the flexibility to accommodate complex designs. Approaches utilizing generalized additive models (GAMs) [11, 12], including tradeSeq [5] and PseudotimeDE [8], are difficult to interpret when compared to traditional linear models, despite their ability to account for the nonlinearity of gene dynamics. Specifically, GAMs utilize knots (locations along pseudotime) to fit smooth functions; however, existing approaches pre-specify knot locations or constrain all genes to have the same knots, limiting their potential biological relevance.

The scGTM method [10] attempted to address the issue of interpretability by classifying dynamic genes into three broad patterns: monotone, hill-shaped, and valley-shaped. However, predefined categories still limit between-lineage comparisons to visual comparison [10]. Similarly, the switchDE methodology [13] aimed to improve biological interpretability by modeling genes using a sigmoid function over pseudotime, allowing the user to identify points in pseudotime at which genes switch from being upregulated to downregulated. However, switchDE models gene expression using a Gaussian distribution, which is not appropriate for scRNA-seq data [14–18], and does not include functionality to account for multiple lineages.

Most current approaches are also unable to account for population structure in datasets composed of cells from multiple subjects, which are becoming more common in properly designed and powered experiments [1]. The issue of multi-subject data was considered in the recent development of Lamian [6], which utilizes subject-level random effects to control for variability between subjects. While this helps to control the false discovery rate, subject-specific inference is not provided, and the use of natural cubic splines limits interpretability and comparability of expression patterns. In addition, Lamian models depth-normalized and log-transformed expression using a Gaussian distribution for the sake of computational efficiency, despite the non-Gaussian nature of scRNA-seq data [14–18]. Finally, Lamian suffers from one of the same drawbacks as tradeSeq - though Lamian does estimate the optimal number of knots in a gene-specific manner, it places them at evenly-spaced quantiles, and thus the knots themselves lack biological interpretability.

To fill this methodological gap, we present scLANE, an interpretable framework for modeling nonlinear expression dynamics and performing trajectory differential expression analysis (**Fig. 1**). scLANE handles a variety of experimental designs inherent to multi-sample scRNA-seq experiments, and can be used downstream of any pseudotemporal or RNA velocity estimation method - making it easy to add to existing analysis pipelines. For each gene within each lineage, scLANE characterizes expression dynamics by identifying locations of expression shifts along the trajectory and quantifying the change in gene expression over these pseudotime intervals. In multi-subject designs, scLANE provides both population-level and subject-specific effect estimates. We demonstrate scLANE’s performance in terms of accuracy and computing efficiency in simulation studies and four case-studies spanning a variety of biologically interesting scenarios. scLANE is freely available and accessible via a web server leveraging high performance computing resources at https://sclane.rc.ufl.edu/. It is also available on Bioconductor (in-process) and GitHub at https://github.com/jr-leary7/scLANE.

## 2 Materials and Methods

### 2.1 Modeling expression over pseudotime: an overview

Without loss of generality, we describe the scLANE model for a single gene and a single lineage. Let *y* be a vector of the observed expression values and *t* be the vector of estimated pseudotime. To motivate scLANE intuitively, we first explain our approach in a general sense before adapting it to the unique statistical challenges of single-cell RNA-seq data.

We begin by assuming that expression counts may be modeled as a function of pseudotime like so, where *f*(·) is some nonlinear function:

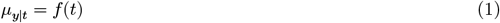

For scLANE, we propose that we can accurately characterize the relationship between pseudotime and gene expression by approximating *f*(·) as a piecewise linear regression model, cast as a set of *p* simpler functions defined over specific subregions along *t* such that:

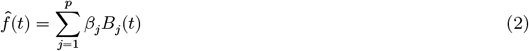

where *B*_*j*_(*t*) are basis functions which span the entire region and are defined so as to achieve continuity. In scLANE we utilize truncated power basis functions, defined as:

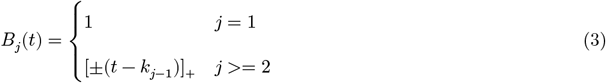

where [*z*]_+_ = *z* if *z* > 0. Thus, the basis functions specify subregions along *t* joined at points *k*, which are referred to as knots. Subregions will indicate changes in expression across the trajectory with the knots representing the timing of expression changes relative to pseudotime. The coefficients [*β*_1_, …, *β*_*j*_] are directly interpretable as multiplicative effect sizes due to their estimation via generalized linear models [20].

**Figure 1.**
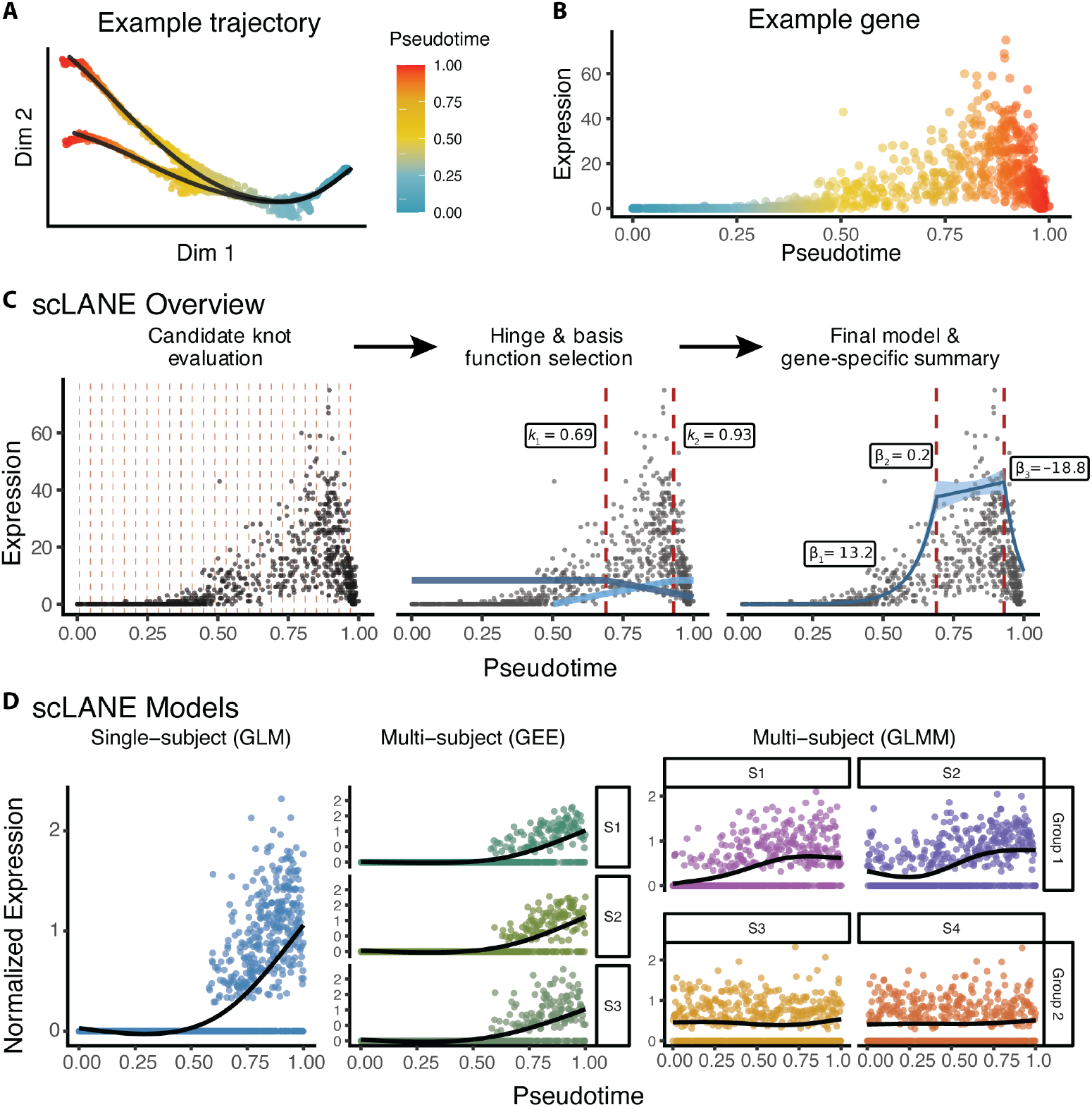
Overview of the scLANE trajectory DE workflow. **(A)** Pseudotime estimates in dimension-reduced space for the zebrafish dataset [19]. Principal curves representing the two-lineage trajectory are overlaid in black. **(B)** Scatterplot of pseudotime versus gene expression for a single dynamic gene. **(C)** Overview of the scLANE modeling procedure. First, candidate knots (dashed lines in red) are evaluated and scored. Once the optimal number and location of knots are selected, the left and right hinge functions at each knot are evaluated sequentially. Gene expression is then modeled as a linear combination of hinge functions, and the final scLANE model is compared to a null model. Model outputs include fitted values with 95% C.I.s, knot locations (*k*), and coefficients for each segment (*β*). **(D)** Simulated gene expression dynamics demonstrating the use cases for the GLM, GEE, and GLMM modes of scLANE, from left to right. The prefix “S” denotes differing samples/subjects.

In order to estimate an optimal partitioning of the pseudotime space into subregions that accurately characterizes a gene’s dynamics, scLANE adopts the multivariate adaptive regression splines (MARS) approach [21]. Briefly, the strategy follows a stepwise regression procedure, where the forward stage partitions the pseudotime space into a specific number of optimal subregions by minimizing a lack-of-fit function. A backward selection stage then prunes the subregions to prevent overfitting. This framework was extended to generalized linear models and generalized estimating equation frameworks by modifying both the lack-of-fit criteria and the backward pruning step and was denoted MARGE [22]. We further extend the MARGE approach for single-cell data. Specifically, we define the following:

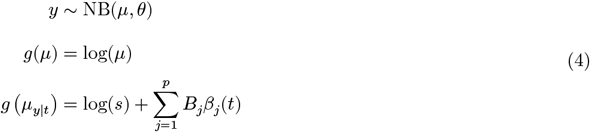

where *μ* is mean expression, *θ* is the overdispersion parameter, and *s* is an offset to account for technical variability in sequencing depth. By default, we formulate the offset by multiplying the depth-normalized counts of each cell by 10,000 to match the counts-per-10k normalization used by the `Seurat` [23] and `Scanpy` [24] packages, though users may easily specify their own offset.

The setup described above covers the single-subject case, which in scLANE is referred to as the GLM mode. We will first describe each step in more detail for this mode, followed by details on the two multi-subject modes below and additional details shown in **Suppl. Table 1**. An overview of the mode-agnostic steps in scLANE testing are presented in **Algorithm 1**. The required inputs are a matrix of unnormalized expression counts, a matrix of pseudotime values with cells as rows and columns as lineages, a vector of subject IDs if cells were taken from multiple subjects, and a sequencing depth offset (if desired).

In order to reduce runtime, model fitting is parallelized across genes using the `foreach` [25] and `doSNOW` [26] R packages. Memory usage is kept to a minimum by storing the counts matrix as a read-only temporary file [27]. In addition, matrix operations such as multiplication and inversion are optimized via the `Rcpp` [28] and `RcppEigen` [29] R packages.

#### Algorithm 1

scLANE testing

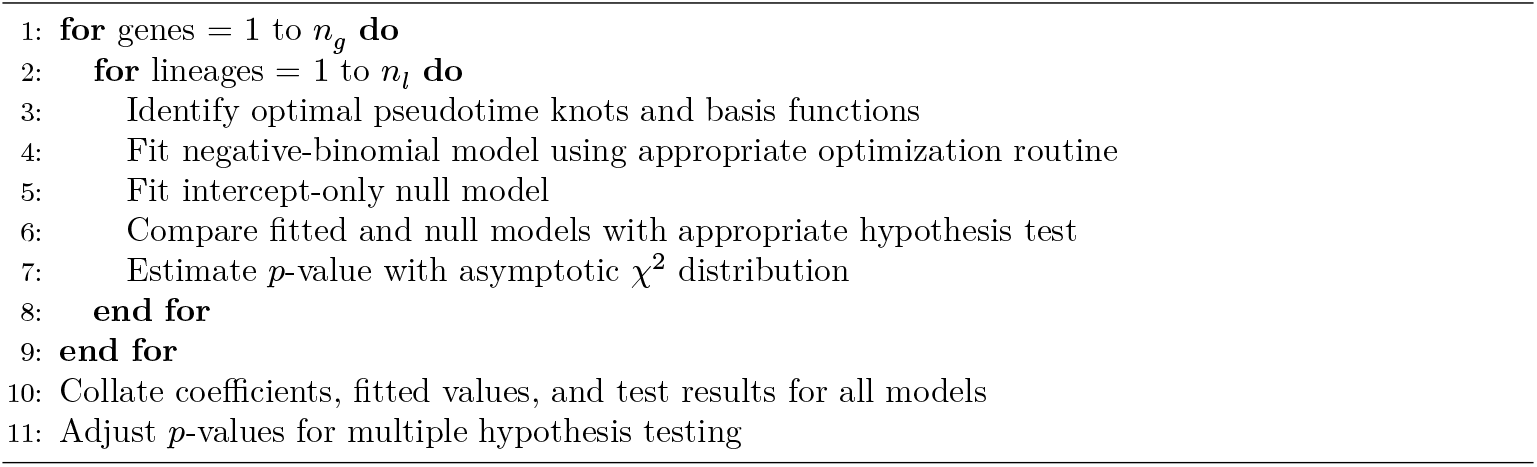

#### 2.1.1 Single subject - GLM mode

For single-cell data, we expect few knots will be chosen in practice, thus we gain computational efficiency by reducing the set of candidate knots from the total number of unique pseudotime values to a default of *n*_*k*_= 25 evenly distributed across the bounds of pseudotime. We also remove candidate knots that are close to the upper or lower bound of pseudotime and require heuristically-estimated minimum and maximum distances between candidate knots [22]. The forward selection stage identifies optimal knots from these candidates using a score statistic as the lack-of-fit criterion, up to a maximum possible number of knots and their associated basis functions, *M* = 3 by default. The backward stage prevents overfitting, with a Wald information criterion being used to prune the model [22]. The final model coefficients are estimated via iteratively-reweighted least squares [20, 30]. For the GLM mode, a negative-binomial model with the canonical log link function is fit using the MASS R package [31]. A likelihood ratio test (LRT) is used for hypothesis testing, where the test statistic is assumed to follow a 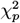 distribution with *p* degrees of freedom. The null model is an intercept-only model matching the structure of the scLANE model. After all models have been fit, *p*-values are adjusted using the Benjamini and Hochberg FDR correction [32].

#### 2.1.2 Multiple subjects - GEE mode

For studies with multiple subjects in which gene dynamics can reasonably be assumed to be conserved across subjects, we developed an extension to the GLM mode based on estimating equations. While the MARGE method extended the classical MARS algorithm to the multi-subject domain, it did not support negative-binomial GEEs [22]. The architecture of the GEE mode is similar to that of the GLM mode, with the estimating function given as follows, where *N* is the total number of subjects, *ϕ* is a scale parameter that is estimated from the data, and *α* is the estimated correlation parameter [33]:

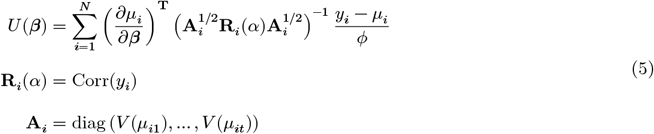

The estimating function is optimized via Fisher scoring [22, 33, 34]. While multiple correlation structures are supported in scLANE, the default is the autoregressive (AR-1) structure, which is preferred when observations that are closer in time can be assumed to be more highly correlated [33, 35]. In our case, since pseudotime is driven by transcriptomic similarity it follows that observations near one another in pseudotime will have correlated measurements of expression. Model fitting is then performed via the geeM R package [34]. The null model is fit using the same correlation structure as the full model, and a Wald or Score test - depending on user preference - is performed to test for significance [34, 36]. Finally, *p*-values are estimated and adjusted as described for the GLM mode.

#### 2.1.3 Multiple subjects - GLMM mode

Lastly, when subjects are taken from multiple groups or it can be assumed that gene dynamics will differ in meaningful ways between subjects, we turn to mixed models to estimate both population-average and subject-specific trends. The general form of the GLMM framework is shown below, with **B** being the matrix of basis functions, and **Z** the matrix of subject-level random effects.

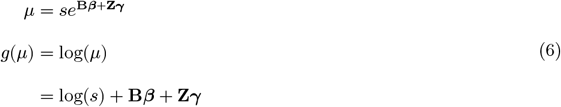

To ensure the model accommodates subject-level expression dynamics, the GLM mode model is fit per-subject with a maximum of *M* = 3 optimal knots as described previously. The basis functions are then recomputed over all subjects simultaneously, and an additional pruning step is performed to ensure that the chosen per-subject basis functions are not highly collinear. This last pruning stage fits a set of negative-binomial LASSO models [37] with cross validation to select the penalty parameter *λ* using the `mpath` R package [38]. The model with the lowest Bayes Information Criterion (BIC) value is considered optimal, and the basis functions with non-zero coefficients are retained. The final GLMM is obtained by fitting the selected basis functions including a random intercept for each subject and per-subject random slopes on each basis function [39] using the `glmmTMB` R package [40]. A LRT is performed between the final and null model. The null model here includes a random intercept for each subject in order to account for differing levels of expression across subjects. The test statistic is assumed to follow an asymptotic 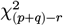 distribution, where *p* is the number of final basis functions, *q* is the number of total random effects, and *r* is the number of random intercepts. Estimation and adjustment of *p*-values is then performed as described for the GLM and GEE modes.

### 2.2 Simulated datasets

To obtain unbiased estimates of trajectory DE method performance, we simulated 72 single-subject and 715 multi-subject UMI-based scRNA-seq datasets under three different parameter regimes via the `Scaffold`R package [41] (**Suppl. Tables 2-4**). The datasets fall into three broad categories: cells from a single subject, cells from multiple subjects with gene dynamics largely conserved between subjects, and cells from multiple subjects with heterogeneous gene dynamics. These three regimes represent the use cases for the GLM, GEE, and GLMM modes implemented in scLANE. Under each regime, we generated datasets using three different reference datasets; the first is composed of healthy human pancreas cells [42], the second of human embryonic neurons [43], and the last of healthy murine pancreatic cells undergoing endocrinogenesis [4]. Prior to simulation, each reference dataset was filtered to ensure that all genes were present in at least three cells. The simulation method implemented in the `Scaffold` R package ensured that regardless of which reference was used, all simulated datasets reflected the sparsity and overdispersion inherent to scRNA-seq count data [16, 41, 44, 45] (**Suppl. Fig. 1**). For single-subject datasets we varied the number of cells generated from 100 − 5000 and the percentage of truly dynamic genes from 1% − 20%. In addition to those parameters, for multi-subject datasets we also varied the sample allocation between subjects (balanced vs. unbalanced) and to what degree gene dynamics were conserved at the population (40%, 50%, 60%) and sample-group levels (70%, 80%, 90%). Each simulated dataset was then preprocessed with a typical workflow [46] built with the `Scater` [47] and `Scran` [48] R packages. For each simulated dataset we ran scLANE along with several other competing methods using their recommended or default settings. Specifically, for single-subject simulations we compared scLANE to tradeSeq [5] and PseudotimeDE [8], and for multi-subject simulations we compared scLANE to tradeSeq and Lamian [6]. Reproducibility was ensured and stochasticity controlled through the implementation of the above as a pipeline using the `targets` R package [49].

For scLANE, a gene was denoted as trajectory DE if its adjusted *p*-value was below *α* = 0.05 (for at least one subject in GLMM mode). We then computed a suite of classification performance metrics for the predicted versus true gene statuses including accuracy, sensitivity, specificity, AUC-ROC, F-measure, and balanced accuracy, the formulae for which are shown in Equation 7. T and F stand for true and false, and N and P for negative and positive, respectively.

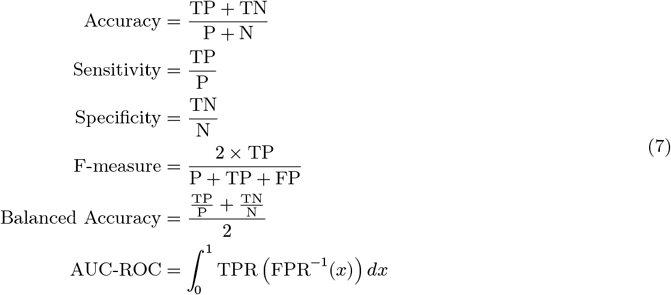

Unless otherwise specified, all methods were run on a high performance computing node with 4 threads used to parallelize model fitting. Specifically, due to its relatively computationally intensive approach, PseudotimeDE was run with 24 threads.

### 2.3 Case studies

Below we describe the data processing for four case-study datasets used to demonstrate scLANE’s performance in real-world scenarios.

#### 2.3.1 Pancreatic endocrinogenesis dataset

The raw spliced and unspliced mRNA counts were obtained from the `scVelo` Python package [50]. We filtered out cells with a sequencing depth of less than 1,000, and genes with fewer than five spliced mRNA counts. Next, 3,000 HVGs were selected using the Seurat method of variance estimation [23, 24]. The counts were then log1p-transformed and scaled prior to linear embedding with PCA [51, 52], after which 30 components were retained and used to create an SNN graph with parameter *k* = 20 and the cosine distance metric [24]. Smoothed first-order moments of the spliced and unspliced counts were estimated using the set of nearest neighbors for each cell [50]. The `scVelo` Python package was used to estimate RNA velocities with the dynamical model, followed by an estimate of gene-shared latent time ordering [50]. Additional details on obtaining the latent time ordering for this dataset is provided in the **Supplementary Methods**.

scLANE was run in GLM mode using the latent time ordering and the per-gene spliced mRNA counts for 4,000 candidate genes having high mean and standard deviation (SD) and low sparsity. tradeSeq [5] was run on the dataset using the same computational resources as scLANE. The optimal number of knots was chosen to be *k* = 5 by evaluating *k* = 3 − 10 and selecting the *k* with the lowest mean absolute deviation from the median AIC as described in the tradeSeq vignette.

A gene regulatory network (GRN) was also estimated on normalized spliced mRNA expression using the random forest (RF) method implemented in the `GENIE3` R package [53]. We tested only the set of 4,000 candidate genes from scLANE against a set of 117 previously-identified murine transcription factors (TFs) [54]. The set of weighted links between TFs and putative target genes, were used to further validate the relationships between regulatory and target genes identified by scLANE.

#### 2.3.2 Zebrafish dataset

The zebrafish dataset [19] was downloaded using the `CellRank` Python package [55]. The original force-directed graph embedding was retained, and after filtering out cells with null embedding values and genes expressed in less than five cells the dataset comprised 15,363 genes across 2,341 cells. The cells were taken from 12 timepoints between 3.3 and 12 hours post-fertilization [19]. After counts-per-10k depth normalization and log1p-transformation, 3,000 HVGs were selected and scaled using `Seurat` [23]. PCA [51, 52] was used to reduce dimensionality, after which an SNN graph with *k* = 20 neighbors was computed on the first 30 PCs using the cosine distance. Lastly, cells were partitioned into 8 clusters via the Leiden algorithm [56] with resolution set to *r* = 0.5. Pseudotime was estimated for two lineages with the `Slingshot` R package [57] based on the first two force-directed graph components, with cluster five set as the root node based on its being composed of early blastomere cells from the earliest timepoint.

scLANE testing was performed for each lineage with default settings on the top 4,000 candidate genes, and *p*-values were adjusted via the Benjamini-Hochberg FDR correction [32]. tradeSeq [5] was run as described previously with *k* = 5 knots using the same computational resources as scLANE. Finally, we used the `patternTest()` function in tradeSeq to identify genes that exhibited significantly different dynamics between the two lineages.

#### 2.3.3 Fetal liver hematopoiesis dataset

The hematopoiesis dataset was downloaded from the Human Developmental Cell Atlas portal [58] and subset the cells to just the six celltypes relevant to the Kupffer cell lineage using the authors’ original annotations [59]. All preprocessing steps were performed using the `Scanpy` Python package [24]. After selecting 4,000 HVGs, we trained an scVI model [60] for 150 epochs with two layers and a negative-binomial likelihood to integrate samples across the fetal ID from which cells were taken. Gene dispersions were estimated within each batch and celltype. The scVI model provided a 20-dimensional latent representation of the cells which was then used as a drop-in replacement for the usual PCA embedding. Using `Scanpy` [24], a nearest neighbors graph was estimated in the latent space with *k* = 20 using the cosine distance. The SNN graph was used as the initialization for a force-directed graph embedding computed using the ForceAtlas2 algorithm [61]. After estimating pseudotime over a single lineage with Slingshot [57], trajectory DE testing was performed with scLANE’s GLM mode on the set of 4,000 candidate genes. We also performed trajectory DE testing with tradeSeq [5] using *k* = 5 knots.

#### 2.3.4 B-cell dataset

We downloaded the human fetal liver hematopoiesis dataset [59] from the Human Developmental Cell Atlas portal [58], then subset the data to just the B-cell lineage. Using the `Scanpy` package [24] we filtered out cells with a sequencing depth of less than 1,000 or a percentage mitochondrial reads greater than 15%, and genes expressed in fewer than five cells. Next, we identified 4,000 HVGs using the “seurat_v3” method, then performed celltype-aware integration via a variational autoencoder (VAE) trained for 150 epochs using the `scVI` package [60]. We then depth- and log1p-normalized the counts, scaled them, and generated a 50-dimensional PCA embedding [51, 52]. After estimating an SNN graph in the VAE latent space with *k* = 50, we clustered the cells with the

Leiden algorithm [56]; in addition we estimated smoothed first-order moments of the counts across each cell’s 30 NNs using the `scVelo` package [50]. We then generated a diffusion map [62] embeddings in two dimensions and estimated a diffusion pseudotime ordering for the cells using the first diffusion component [62].

We performed subject-aware scLANE testing using the GEE mode after identifying 4,000 candidate genes. We also performed subject-naive trajectory DE testing via tradeSeq [5], along with subject-aware testing using Lamian [6]. Geneset enrichment was performed using the gprofiler2 R package [63].

## 3 Results

### 3.1 Simulation study

To first validate scLANE’s ability to accurately classify genes as dynamic or static, we generated a total of 787 simulated datasets under three different experimental design setups (**Suppl. Fig. 1**). For single-subject simulations, we compared the GLM mode of scLANE to tradeSeq [5] and PseudotimeDE [8]. Multi-subject simulations were split into two categories: those with gene dynamics mostly conserved between subjects and those with multiple heterogeneous subject groups. These two setups correspond to the use cases recommended for the GEE and GLMM modes of scLANE, respectively. We compared the GEE and GLMM modes of scLANE to Lamian [6].

On single-subject simulated datasets, scLANE compared favorably to tradeSeq and PseudotimeDE, exhibiting higher or roughly equal performance on classification metrics such as balanced accuracy, F-measure, sensitivity, and specificity **(Fig. 2A, Suppl. Fig. 2A-C)**. This trend held across varying dataset sizes, reference datasets, and dynamic gene frequencies, where scLANE classified truly dynamic and truly static genes more accurately than the other methods (**Suppl. Fig. 2D**). In addition, memory usage was significantly and consistently lower for scLANE across all simulations (**Fig. 2B**). The largest datasets we simulated were composed of 5,000 cells; on those datasets, scLANE used an average of just 0.427 gigabytes of memory. With respect to runtime, scLANE also processed more genes per minute on average, and on the largest datasets, scLANE, tradeSeq, and PseudotimeDE processed 7.1, 5.9, and 1.1 genes per minute on average, respectively **(Fig. 2C)**.

scLANE also out-performed other approaches for data simulated with multiple subjects. For GEE mode, scLANE had a consistently larger or equivalent balanced accuracy compared to Lamian (**Fig. 2D**). As expected, datasets with larger numbers of cells resulted in better classification of truly dynamic genes for both methods. Lamian struggled more on datasets composed of fewer than 1,000 cells, while scLANE was still able to achieve high classification accuracy. For instance, on datasets composed of 1,000 cells, scLANE exhibited a mean balanced accuracy of 0.949, while Lamian had a mean of 0.872.

The same was true for multi-subject datasets where trajectories were less conserved between subjects for evaluating scLANE’s GLMM mode performance (**Fig. 2E**). In general, scLANE-GLMM was the least affected by differences in the dynamic overlap due to its subject-specific nature. In comparing each method’s performance between scenarios with relatively high (90%) versus low (70%) dynamic gene overlap among subjects, scLANE-GLMM had only an average 2.4% loss in balanced accuracy, while the GEE mode and Lamian had a 5.3% and 5.0% loss, respectively. We observed that scLANE’s GEE and GLMM mode used more memory than Lamian, although all three methods were memory efficient using less than 0.5 gigabytes in every scenario (**Suppl. Fig. 2E**). scLANE’s GEE mode was consistently faster than Lamian in processing time, whereas the GLMM mode was slightly slower for larger datasets (**Suppl. Fig. 2F**).

### 3.2 Case study results

Next, we demonstrate scLANE’s ability to generate interpretable biological insights on four published datasets: murine pancreatic endocrine cell fate specification, zebrafish axial mesoderm development, human fetal liver hematopoiesis, and human B-cell maturation. Processing details are described in **Methods**.

#### 3.2.1 Murine pancreatic endocrinogenesis

We applied scLANE to a well-annotated dataset composed of 3,696 *Mus musculus* cells during pancreatic endocrinogenesis (**Fig. 3A-B, Suppl. Fig. 3A-B**) [4]. While this dataset has been widely used in benchmarking trajectory methods and package vignettes, the biological properties of the dataset have often been left unexplored beyond recapitulating the original analysis, presenting an opportunity to showcase the biological utility of scLANE. After estimating an RNA velocity-based latent time [50], scLANE identified a set of 3,355 trajectory DE genes using the GLM mode. scLANE identified relevant dynamics in the genes producing cell-type specific peptide in the four mature endocrine cell types (**Suppl. Fig 3C**), and the distribution of knots across all genes showed many early knots in the progenitor stage (**Suppl. Fig 3D**). The early knots indicate that specification occurs prior to the appearance of the mature endocrine cells.

We first examined the dynamics of transcription factors (TFs) known to regulate pancreatic endocrine cell differentiation, specifically: *Sox9, Neurog3, Rfx6, Arx, Pax6, Pax4, Pdx1*, and *Neurod2* [64–69] (**Fig. 1C-E**). *Neurog3* is the master regulator and driver of pancreatic endocrinogenesis, whereby it guides specification of alpha and beta cells during the pre-endocrine stage [64]; it is also regulated by *Sox9* expression [70]. We observed this in the dynamics of both genes, where *Sox9* is active at the beginning of latent time, appearing to activate *Neurog3*, which continued increasing in expression until midway through the trajectory (knot = 0.454), then decreasing as the alpha and beta cells furthered in maturity (**Fig. 3C**). While *Neurog3* initiates differentiation, the commitment to the alpha cell phenotype is also partially driven by expression of *Rfx6, Arx*, and *Pax6* [65, 66], all of which experienced peak expression near (in latent time) to where the alpha cell population was located **(Fig. 3D)**.

The peak expression knot for *Neurog3* was estimated by scLANE to be earlier than its validated target – *Arx* (knot = 0.856) [67].

scLANE fitted dynamics also help to explain beta cell development. Notably, the upregulation of *Arx* is known to suppress expression of *Pax4* [67], which is observed in the scLANE dynamics of *Arx*, having an extremely swift downregulation (slope = -60.679) occurring near to the appearance of a large group of beta cells (**Fig. 3D-E**). Additionally, *Neurod2* has been shown to be activated *in vivo* by *Neurog3* [68] and linked to the development of beta cells [4]. scLANE placed a peak expression for *Neurod2* immediately prior to the emergence of beta cells (knot = 0.577) (**Fig. 3E**). Similarly, scLANE estimated a knot in latent time prior to the appearance of any beta cells (knot = 0.33) for *Pdx1*, where its expression switched from being relatively static to increasing as reflected in the estimated scLANE slope coefficients before (−1.755) and after (2.682) the knot (**Fig. 3E**). *Pdx1* directly targets the insulin gene *Ins1* [64, 69] - insulin being the primary peptide product of beta cells.

The regulation of alpha versus beta cell fate specification and corresponding competing expression of *Arx* and *Pax4* is still not completely understood. We next sought to identify genes with similar dynamics to *Neurog3*, which activates both TFs [64]. After embedding the estimated scLANE gene dynamics in a KNN graph, we identified five genes most similar to *Neurog3*, all of which had peak expression around the same location in latent time (mean knot = 0.478) (**Fig. 3F**). The third-most similar gene was the cell cycle inhibitor *Gadd45a*, previously identified as upregulated in endocrine progenitors and associated with endocrine cell fate specification [4]. However, the most-similar gene, *Btbd17*, has no known function in endocrinogenesis - though it has been shown to be differentially expressed in endocrine progenitor cells [65]. Finally, *Cbfa2t3* has not been previously shown to play a role in endocrine cell fate specification, but is a well-validated transcriptional co-repressor during hematopoiesis, and also promotes myeloid differentiation during acute myeloid leukemia [71–73].

To validate the regulatory potential of these candidate genes, we estimated a gene regulatory network and compared the top regulators of *Btbd17, Gadd45a*, and *Cbfa2t3*. For all three genes, the most highly-weighted TF was the master regulator *Neurog3*, with weights of 0.164, 0.147, and 0.135, respectively. All three links were within the top 23 links out of a total of 705,775 nonzero links identified by the network.

**Figure 3.**
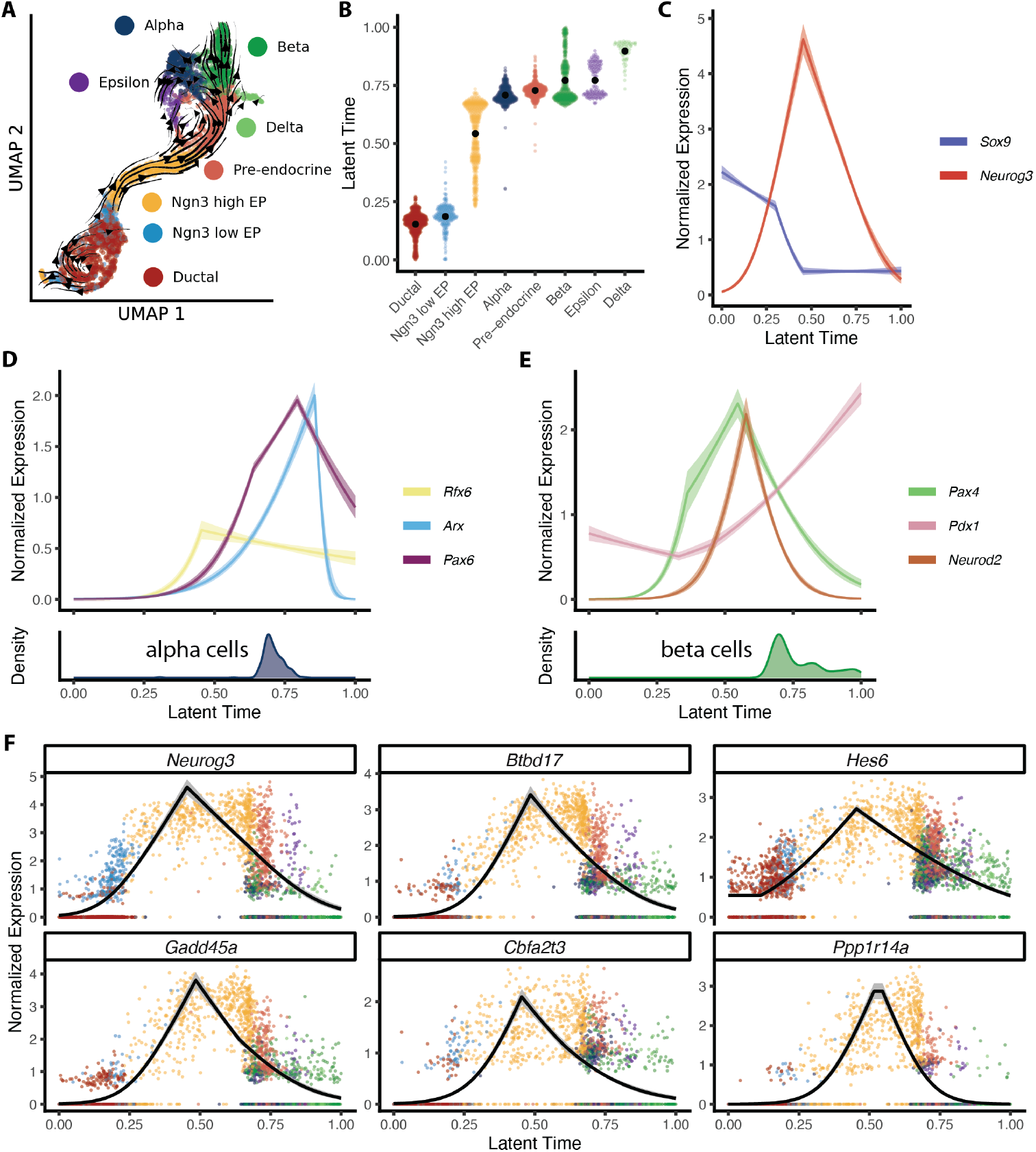
scLANE identifies biologically meaningful and specific gene programs regulating endocrinogenesis. **(A)** Streamline plot of differentiation direction vectors projected onto the UMAP embedding. **(B)** Beeswarm plot displaying the distribution of latent time estimates within each celltype. The mean value per-celltype is displayed in black. **(C)** Fitted dynamics of TFs that act as master regulators of endocrine differentiation. **(D)** Fitted dynamics of genes regulating alpha cell development across latent time, with the distribution of alpha cells in latent time shown below. **(E)** Fitted dynamics of genes regulating beta cell development as in **(D). (F)** Fitted dynamics of the five nearest-neighbors of *Neurog3* across latent time.

#### 3.2.2 Zebrafish musculoskeletal system development

We next applied scLANE to a temporally-resolved zebrafish embryogenesis dataset [19] to demonstrate its utility in analyzing a multi-lineage topology. We chose to focus on characterizing the early dynamics of axial mesoderm differentiation into the fates of the notochord and prechordal plate. We estimated the trajectory and pseudotimes using Slingshot [19, 57] (**Fig. 4A-B, Suppl. Fig. 4**). Pseudotime was rescaled to a common range (**Supplemental Methods**) and scLANE was run in default mode to identify trajectory DE genes for both the notochord and prechordal plate lineages. Building lineage-specific models for the top 4,000 candidate genes on 2,341 cells with 16 threads took 12.2 minutes. In comparison, running tradeSeq with the same number of threads took 25.3 minutes.

Overall, the vast majority of the scLANE trajectory DE genes displayed a concave dynamic in both lineages, with 68% having non-decreasing initial dynamics and 75% had non-increasing terminal dynamics across both lineages. Using a ratio of scLANE’s lineage-specific test statistics, we identified genes that exhibited markedly similar dynamics across lineages and those uniquely dynamic to a single lineage. Around 73% of trajectory DE genes were similarly dynamic across the two lineages, having ratios close to one, such as *eef1a1l1* and *rplp1* (**Fig. 4C**). The rest of the genes were approximately evenly divided between being specific to the prechordal plate or the notochord lineage. Genes that exhibited dynamics specific to the prechordal plate lineage included *ctslb* and *lgals3l* and those specific to the notochord lineage included *kdelr2a* and *plp2* (**Fig. 4C**). While *ctslb* is a well-known prechordal marker gene, the roles of *lgals3l, kdelr2a*, and *plp2* in axial mesoderm are not yet clearly defined.

To further demonstrate scLANE’s interpretability compared to other trajectory DE testing approaches, we used the patternTest function in tradeSeq [5] to identify genes with significantly different dynamics between lineages and compared to using scLANE’s ratios of lineage-specific test statistics. Three of the top six genes by tradeSeq (*id3, apoeb, btg2*) showed differing dynamics between the two lineages, with higher specificity for the prechordal plate (**Fig. 4D**), which was also indicated by scLANE’s ratios. Though the remaining genes in tradeSeq’s top six (*tram1, marcksl1b, calr3b*) appeared to have highly similar gene expression values and dynamics between the two lineages (**Fig. 4D**). The scLANE ratios were all close to one and similar to that of the shared genes *eef1a1l1* and *rplp1* in **Fig. 4C**. Overall, there was little agreement between the tradeSeq’s patternTest statistic and scLANE’s ratio (**Fig. 4E**). While scLANE was able to clearly distinguish moderately and highly lineage-specific genes, none of the four highly lineage-specific genes shown in **Fig. 4C** were ranked within the top 100 by tradeSeq. Additionally, the direction of scLANE’s ratio was able to indicate the lineage-specificity, with larger values indicating the notochord and smaller values indicating prechordal plate.

**Figure 4.**
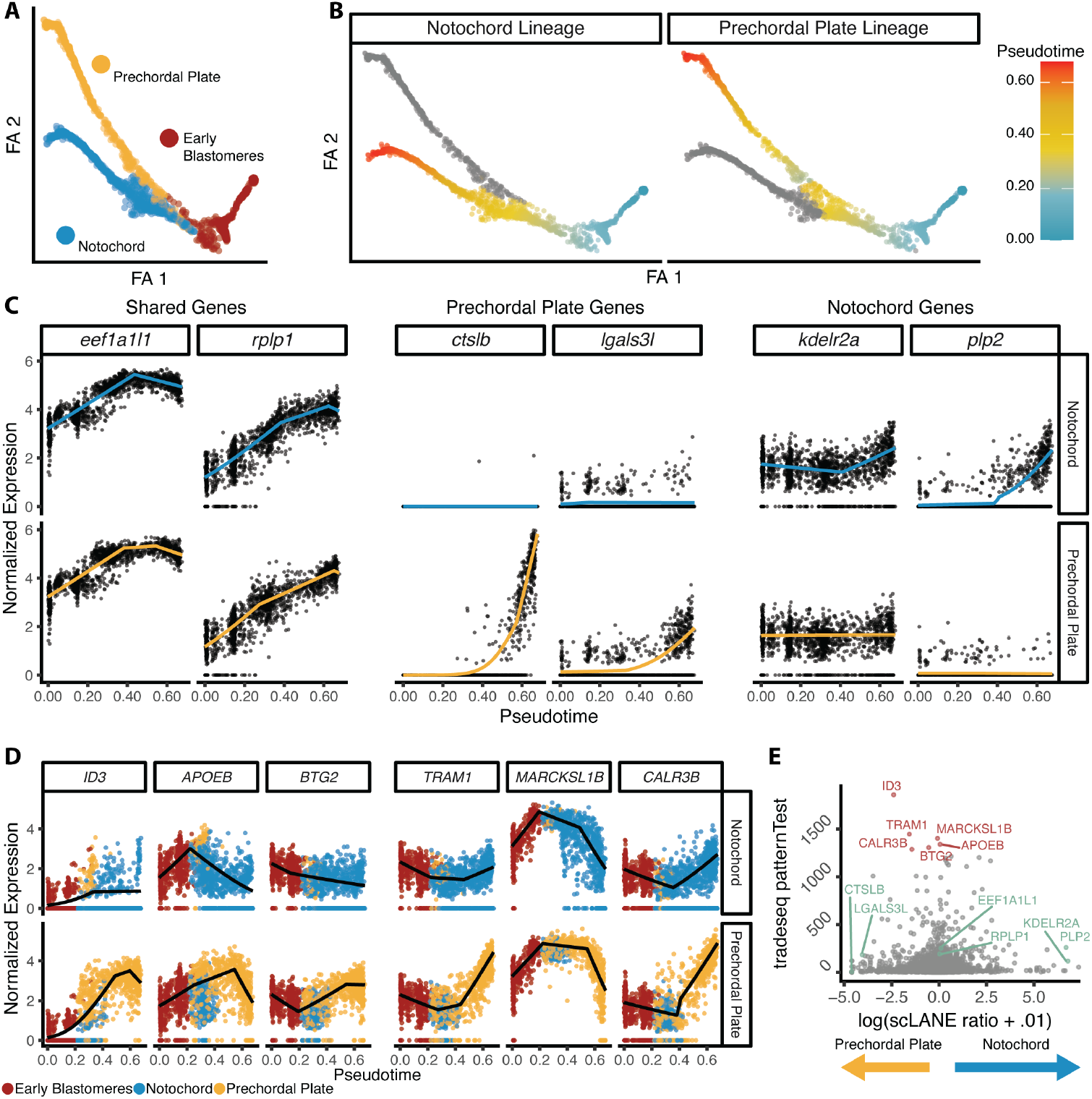
scLANE provides a finer-resolution characterization of zebrafish embryogenesis. **(A)** Force-directed graph embedding of zebrafish axial mesoderm cells colored by celltype. **(B)** Pseudotime estimates for the notochord and prechordal plate lineages. Cells in the prechordal plate were rescaled so that the pseudotime across the two lineages aligned. **(C)** Fitted gene dynamics for selected genes of interest that had trends that were either highly similar across both lineages or highly unique to either the prechordal plate or notochord. **(D)** Scatterplots of gene expression over lineage-specific pseudotime for the top six genes determined to differ most significantly between lineages by tradeSeq’s patternTest function. Fitted dynamics from scLANE are overlaid in black. **(E)** Comparison of scLANE’s ratio of lineage-specific test statistics and tradeSeq’s patterntest for comparing expression between lineages. The six genes of interest identified by scLANE shown in (C) are shown in green and the top six genes by tradeSeq in (D) are colored red.

#### 3.2.3 Scaling up: Monocyte development in human fetal liver cells

Next, in order to showcase the efficiency and scalability of scLANE, we performed trajectory DE testing on a human fetal liver hematopoiesis dataset consisting of 38,188 cells [59]. After preprocessing and pseudotime estimation with `Slingshot` [57], we obtained a single lineage beginning with hematopoietic stem cells and ending in Kupffer cells, with several intermediate monocyte and macrophage phenotypes (**Fig. 5A, Suppl. Fig. 5A-B**). Since human monocytes are relatively well-annotated and their development is largely understood, this dataset served as a confirmatory example of scLANE’s ability to generate valid biological insights at scale. Using the GLM mode of scLANE we performed trajectory DE testing across the sole lineage, and identified 3,325 trajectory DE genes exhibiting highly heterogeneous functionality. The original publication focused on the differences in dynamically-regulated genes between tissues, specifically the dynamics of the canonical monocyte markers *S100A8/A9/A12* and *FCGR1A/2A* [59]. scLANE recapitulated the trajectory dynamics of these key marker genes (**Suppl. Fig. 5C**). However, since their importance to myeloid development and various types of immune response is well-known, we opted to focus dynamics of transcription factors and their regulation of commitment to the macrophage cell fate.

**Figure 5.**
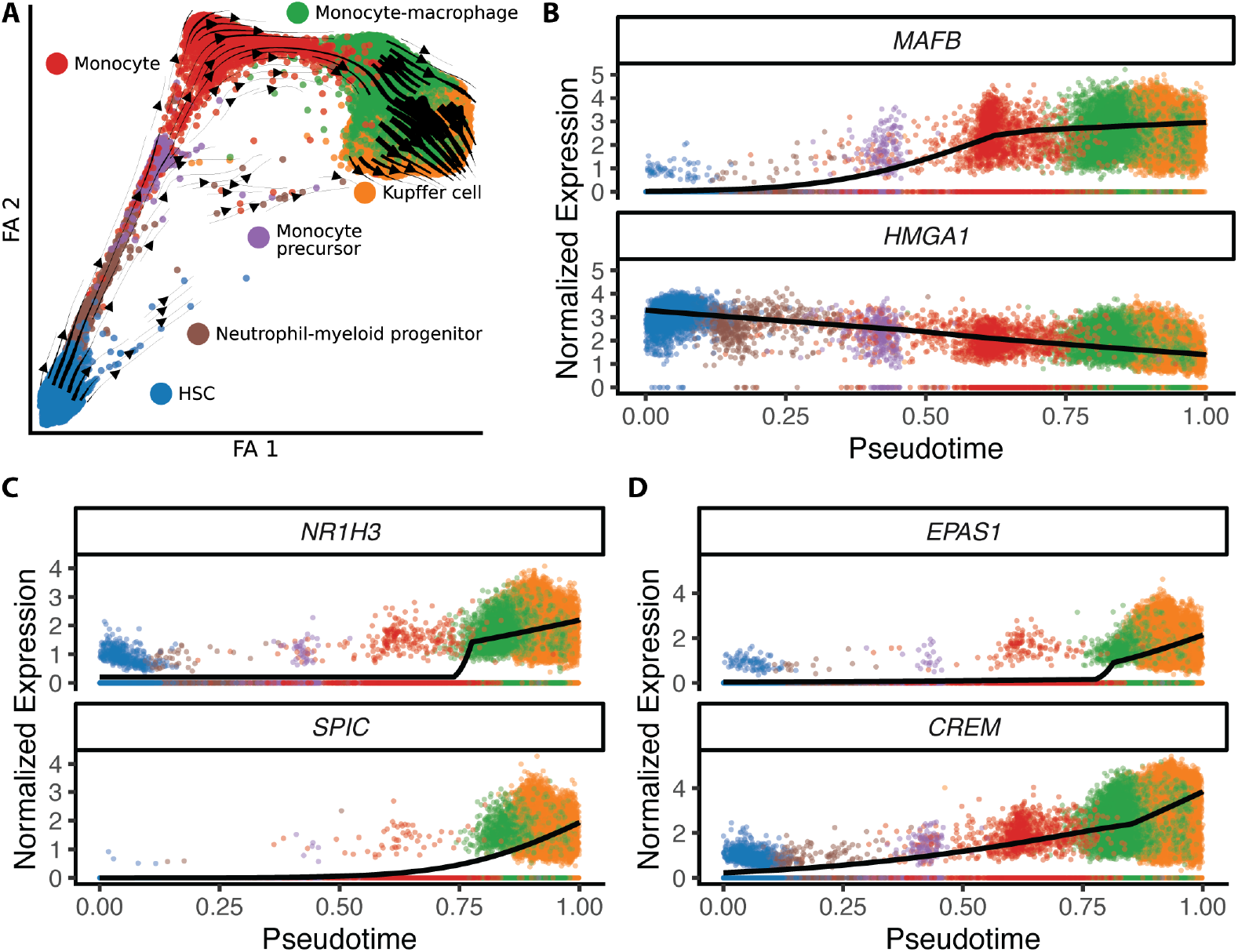
scLANE reveals underexplored TFs at scale. **(A)** Force-directed graph embedding of 38k myeloid cells undergoing hematopoiesis. Differentiation direction vectors are overlaid as streamlines. **(B-D)** Scatterplots of top ranked TFs with dynamics from scLANE overlaid in black.

Ranking genes by their scLANE test statistic, the two highest ranked genes were *MAFB* and *HMGA1* (**Fig. 5B**). *MAFB* is a core macrophage differentiation marker [74] and *HMGA1* plays a prime role in normal hematopoietic differentiation [75]. Within the top six dynamic genes, we also have two Kupffer lineage-determining transcription factors: *NR1H3* and *SPIC* (**Fig. 5C**). These two TFs produce LXR and SpiC which collaboratively bind to open chromatin regions and facilitate Kupffer-specific transcriptional expression and regulation [76]. Two other top TFs, *EPAS1* and *CREM*, also increased in expression along the trajectory toward Kupffer cells (**Fig. 5xsxsD**). *EPAS1* encodes for HIF-2, which was recently found to be differentially regulated in Kupffer cells by Jeelani et al., 2025 [77]. *CREM* has been shown to be involved in directing macrophage cell polarization through its regulation of cAMP-dependent transcription [78] and in Kupffer cells specifically, the cAMP-PKA-STAT3 signaling pathway was shown to direct polarization [79]. With the exception of *SPIC*, the Kupffer specific genes *NR1H3, EPAS1, CREM* all had a knot estimated at 0.78 by scLANE, just prior to the appearance of the Kupffer cells.

#### 3.2.4 Human B-cell maturation

Our final case study concerns the development of B-cells from multiple human fetal livers [59]. After preprocessing the single-lineage dataset, which begins with hematopoietic stem cells before progressing through several progenitor celltypes and terminating in mature B-cells (**Fig. 6A-B**), we performed trajectory DE testing with scLANE, tradeSeq [5], and Lamian [6]. We ran scLANE in GEE mode as the dataset is composed of cells from 14 different subjects with widely varying numbers of cells per-subject (**Fig. 6C**). This process took just 3.6 hours using 24 threads, showcasing scLANE’s ability to efficiently fit and summarize complex multi-subject models. There were a similar number of trajectory DE genes for scLANE and tradeSeq, and a larger set of trajectory DE genes for scLANE than Lamian (**Fig. 6D**).

We compared genes identified as trajectory DE only by scLANE versus those significant only in tradeSeq, and only Lamian. The genes DE by scLANE only had larger coefficients of variation, or variance relative to their mean, whereas genes DE in Lamian only had relatively small variations (**Fig. 6E**). Many of the genes identified by scLANE and not Lamian demonstrated trajectory-relevant variation dynamics, the top four of these genes (ranked by scLANE test statistic) are shown in **Fig. 6F**. We then performed pathway enrichment analysis of the set of genes classified as trajectory DE by scLANE but not by Lamian (**Suppl. Table 5**). Of the terms with adjusted *p*-values less than *α* = 0.1, the most significantly enriched were pathways such as “cell surface receptor signaling pathway” (GO:0007166), “immune system process” (GO:0002376), “regulation of immune system process” (GO:0002682), and “regulation of leukocyte migration” (GO:0002685). All of these terms are clearly relevant to the maturation of human B-cells, indicating that Lamian’s model may have been overly conservative.

**Figure 6.**
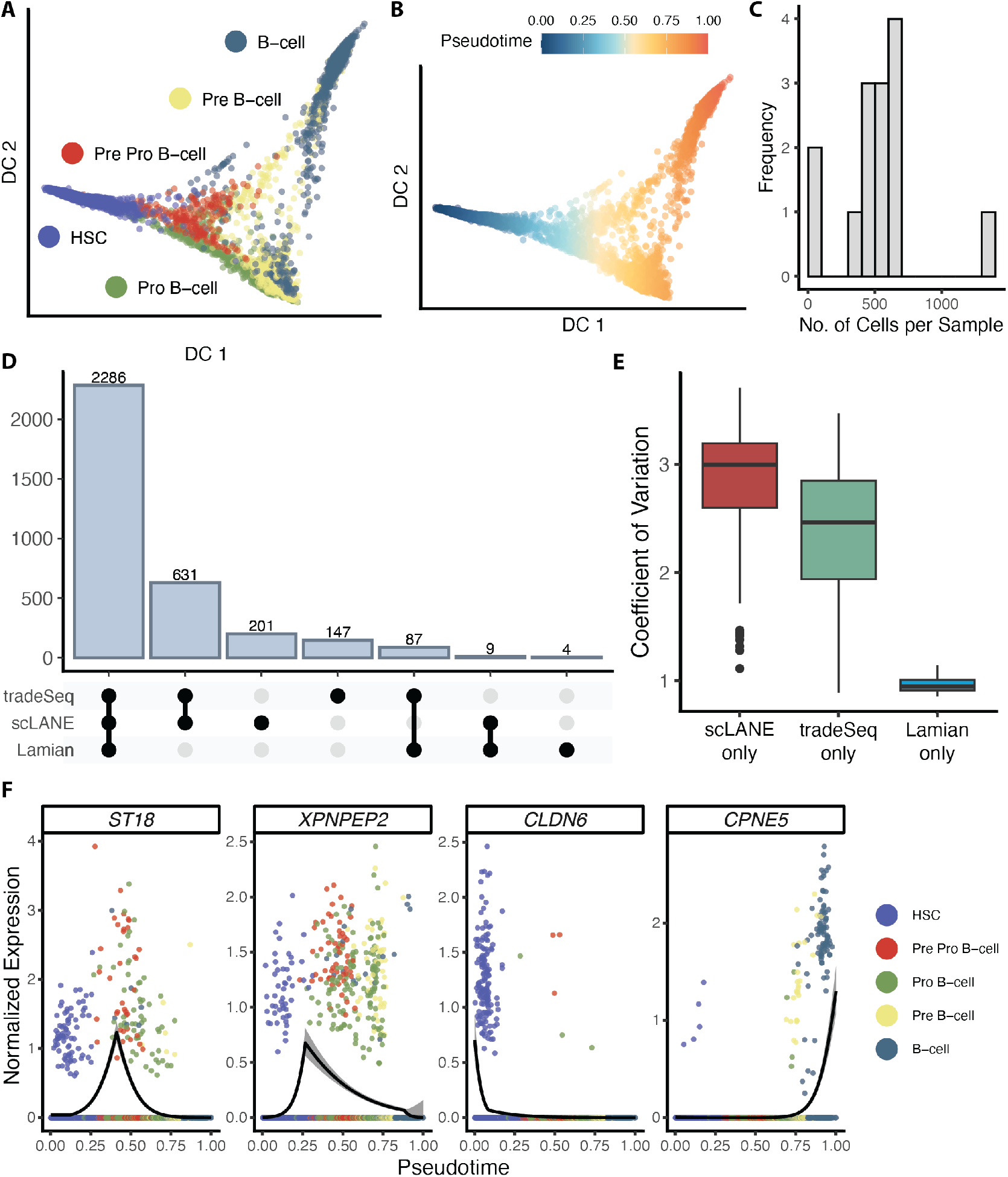
scLANE accurately accounts for multi-subject variability in B-cell maturation. **(A)** Diffusion map embedding of maturing B-cells with points colored by celltype. **(B)** Diffusion map embedding colored by diffusion pseudotime. **(C)** Distribution of the number of cells per sample. **(D)** Upset plot displaying the sets of trajectory DE genes identified by scLANE, tradeSeq, and Lamian. **(E)** The coefficient of variation for genes identified as trajectory DE only by scLANE, only by tradeSeq, and only Lamian. **(F)** Scatterplots displaying gene dynamics for the top genes identified as trajectory DE by scLANE only. The GEE fit from scLANE is overlaid in black along with a 95% C.I. in grey.

## 4 Discussion

A core difficulty in the utilization of scRNA-seq experiments is the complexity of interpreting downstream analysis results. For trajectory analyses, methods too often rely on visual inspection of trends and subjective assessments of whether genes are upregulated, downregulated, or static at any given point in pseudotime. Results are often used to confirm preexisting notions of marker gene behavior. To fully extract novel insights from single cell data, it is imperative that downstream analysis methods carry a high degree of interpretability, in addition to providing quantitative rather than qualitative results. We developed scLANE to address this by providing linear regression-based effect sizes with simple, multiplicative interpretations that any scientist can use to quantify the degree to which genes change during a biological process. Our framework is modular, and can be easily extended to accommodate complex experimental designs while remaining user-friendly. In addition, we have implemented scLANE as a free webserver equipped with high-performance computing resources.

Several challenges remain, in particular with respect to computational efficiency. scRNA-seq datasets have dramatically increased in size during the past five years, and modeling atlas-scale trajectory dynamics is still a time-consuming and resource-heavy endeavor [1, 80]. While we implemented several memory conservation and approximation techniques when developing scLANE, fitting dynamics for larger datasets such as the fetal liver hematopoiesis dataset - composed of almost 40,000 cells - took several hours on a compute cluster when using 16 threads and 150G of memory to test the set of candidate genes. scLANE’s runtime was more demanding in the case-studies compared to existing approaches, however all methods required multiple threads to reduce computational overhead and scLANE offsets its increased time by reducing post-analysis time in providing biologically interpretable results. That said, reducing computation time is an active area of research, and should still be considered a primary concern when improving single cell methods.

Lastly, while we showed scLANE to be remarkably accurate at identifying trajectory DE genes in simulated data, it remains true that in real-world situations the true trajectory is unknown and must be inferred from gene expression data. This presents a challenge due to an additional source of uncertainty, where we stress that care must be taken to validate trajectories biologically prior to performing DE analysis. In every case study,

a number of pseudotime or latent time constructions and embeddings were considered in order to identify one that best recapitulated the underlying biology before proceeding with further analysis. The use of known marker genes is widely used for this process, and recent computational tools may also assist in choosing optimal embeddings [81]. A further challenge in the analysis is the double-dipping issue prevalent in single-cell RNA-seq which can lead to artificially small *p*-values [82]. This is an unresolved challenge in trajectory inference, but may be partially alleviated by using fewer genes to construct the trajectory while testing for trajectory DE on a larger set of genes.

Our extensive simulation study provided a set of benchmarks that point toward scLANE’s ability to accurately and precisely classify genes as dynamic or static under a variety of pseudotime structures while providing quantitative outputs for each gene’s dynamics. In applying our method to a wide variety of developmental datasets, both with respect to species and experimental design, we demonstrated scLANE’s flexibility and ease of relevant biological interpretation. Our four case studies displayed scLANE’s ability to not only recapitulate known biological information by estimating gene trajectory dynamics, but to also perform comparisons across lineages, identify meaningful gene programs, and perform pathway analysis. In our analysis of the endocrinogenesis dataset scLANE identified genes sharing dynamics with the master regulator *Neurog3*, revealing under-explored candidate genes that warrant further study. The multi-lineage zebrafish dataset showcased the ability of scLANE to characterize lineage-specific transcriptional switches controlling cell fate specification. Our investigation of cells undergoing hematopoiesis in the liver demonstrated scLANE’s capability of handing large datasets and generating dynamic gene programs involved in myeloid differentiation, the understanding of which is relevant to the study of immune response. Finally, our analysis of maturing B-cells displayed scLANE’s ability to precisely account for complex experimental designs and control for inter-subject variation when performing trajectory DE testing to extract relevant and interpretable biological signal. Altogether, these results indicate that scLANE was able to accurately characterize the biological mechanisms by which development occurs across multiple different tissues and organisms.

## 5 Declarations

### 5.1 Data & code availability

An R software library implementing the method as described can be downloaded from https://github.com/jr-leary7/scLANE. The pipeline used to generate simulated datasets and benchmark methods is available at https://github.com/jr-leary7/scLANE-Simulations. The code used to generate all results is available at https://github.com/jr-leary7/scLANE-Revision. All processed case study datasets have been deposited on Zenodo at https://doi.org/10.5281/zenodo.10012311.

### 5.4 Author contributions

R.B. and J.R.L. conceived the method and study. J.R.L. and R.B. implemented the software package. R.B. provided statistical guidance and methodological feedback. J.R.L. and X.D. designed the simulation experiments, performed real dataset analyses, and interpreted results. J.R.L. and R.B. wrote and edited the manuscript.

## 5.3 Acknowledgments

We thank Dr. Aaron Molstad for his thoughts on the mixed models implementation. In addition, we thank Drs. Matias Kirst and Wendell Peirera for providing biological motivations and testing the software. Lastly, we thank Jonathan Streater and TJ Schultz for the assistance in creating the scLANE web application.

## 5.4 Funding

This work is supported by National Institutes of Health grant R35GM146895 to R.B.

## 5.5 Conflicts of interest

The authors have no conflicts of interest to declare.

## Notes

https://sclane.rc.ufl.edu/

### Competing Interest Statement

The authors have declared no competing interest.

### Summary of Updates

Revised Methods section and expanded comparisons to other approaches.

