## Supplement for "Interpretable Trajectory Inference with Single Cell Linear Adaptive Negative-binomial Expression (scLANE) Testing"

May 22, 2025

### 1 Supplementary Methods

Briefly, for all datasets, counts were depth-normalized then variance-stabilized using a log1p-transform, after which a set of highly-variable genes (HVGs) was selected and used as input to PCA [1, 2]. The top 30 PCs were then used as initialization for UMAP [3, 4], force-directed graph [5], and diffusion map [6] embeddings in order to embed the cells in two dimensions for visualization purposes. In general, the embedding that best recapitulated the underlying biology was used as the basis for the construction of the trajectory.

#### 1.1 Pancreatic endocrinogenesis dataset

After embedding the cells in 2D via UMAP [3], the RNA velocity vectors were projected onto the embedding and visualized [7]. In addition, when computing the velocity graph we estimated an uncertainty value for each cosine similarity between velocity vectors and true changes in gene expression between cell states. These uncertainty estimates were propagated forward using Monte Carlo sampling to the estimation of a cell-cell transition probability matrix using the velocity kernel implemented in the `CellRank` Python package [8]. Next, CytoTRACE differentiation potential scores and directions were estimated per-cell using the CytoTRACE kernel in `CellRank` [8, 9]. We then used the diffusion pseudotime estimates for each cell to generate a third kernel and accompanying transition probability matrix [6, 8]. Finally, the three kernels were combined with weights of 0.5, 0.25, and 0.25, respectively. The combined kernel's transition probability matrix was projected onto the UMAP embedding and visualized.

#### 1.2 Zebrafish musculoskeletal system development

The two lineages share a common pseudotime before diverging into the prechordal plate and notochord lineages. In order to compare the two lineages' pseudotimes directly, the longer lineage (prechordal plate) was rescaled to have the same maximum pseudotime as the shorter lineage (notochord). The rescaling was done only on the pseudotimes for cells that diverged from the common pseudotime, specifically, cells in the prechordal plate with pseudotime larger than 0.4263415 had the difference between the two lineages max. pseudotime subtracted (0.2086) prior to analysis.

#### 2 Supplementary Tables

| Mode | Null Model Structure | Fitting Method |
|---|---|---|
| GLM | Intercept-only NB GLM | `MASS::glm.nb()` |
| GEE | Intercept-only NB GEE with desired correlation structure | `geeM::geem()` |
| GLMM | NB GLMM with a random intercept per subject | `glmmTMB::glmmTMB()` |

**Supplementary Table 1.** Structures of scLANE's null model for each mode.

| Parameter | Values |
|---|---|
| Number of cells | $100, 250, 500, 1000, 3000, 5000$ |
| Dynamic gene frequency | $0.01, 0.05, 0.1, 0.2$ |

**Supplementary Table 2.** Varying parameters used in the scLANE single-subject (GLM mode) simulation study.

| Parameter | Values |
|---|---|
| Number of cells | $250, 500, 1000, 3000, 5000$ |
| Dynamic gene frequency | $0.05, 0.1$ |
| Dynamic gene overlap frequency | $0.7, 0.8, 0.9$ |
| Subject allocation | Balanced, unbalanced |

**Supplementary Table 3.** Varying parameters used in the scLANE multi-subject (GEE mode) simulation study. Dynamic gene overlap frequency refers to the percentage of dynamic genes whose dynamics were retained between subjects i.e., the simulated knots and slopes were kept the same across all subjects.

| Parameter | Values |
|---|---|
| Number of cells | $250, 500, 1000, 3000, 5000$ |
| Dynamic gene frequency | $0.05, 0.1$ |
| Dynamic gene overlap frequency | $0.7, 0.8, 0.9$ |
| Subject allocation | Balanced, unbalanced |
| Dynamic gene group overlap frequency | $0.4, 0.5, 0.6$ |

**Supplementary Table 4.** Varying parameters used in the scLANE multi-subject, multi-group (GLMM mode) simulation study. Dynamic gene overlap frequency refers to the percentage of genes whose dynamics were retained across subjects within each group (like in the multi-subject GEE mode study), while group overlap frequency refers to the same but across subject groups.

| Term ID | Term Name | Adjusted *p*-value |
|---|---|---|
| GO:0009605 | response to external stimulus | 2.2970724741590474e-11 |
| GO:0002376 | immune system process | 4.1614064915571264e-11 |
| GO:0006955 | immune response | 1.2267249684916839e-10 |
| GO:0051239 | regulation of multicellular organismal process | 1.3893786577134221e-10 |
| GO:0006952 | defense response | 4.0311756496437644e-10 |
| GO:0006950 | response to stress | 4.057191967551892e-10 |
| GO:0007166 | cell surface receptor signaling pathway | 5.654518868750895e-9 |
| GO:0002682 | regulation of immune system process | 6.681954267681861e-9 |
| GO:0006935 | chemotaxis | 2.9171996788336506e-8 |
| GO:0042330 | taxis | 3.3251887892857826e-8 |
| GO:0040011 | locomotion | 1.2036295265290826e-7 |
| GO:0016477 | cell migration | 1.6639981648550187e-7 |
| GO:0048522 | positive regulation of cellular process | 2.1669033192308827e-7 |
| GO:0001568 | blood vessel development | 4.899209531430492e-7 |
| GO:0002685 | regulation of leukocyte migration | 8.180865838695716e-7 |
| GO:0032101 | regulation of response to external stimulus | 1.0838810783188887e-6 |
| GO:0001944 | vasculature development | 1.988074902392259e-6 |
| GO:0098542 | defense response to other organism | 2.0400193921329672e-6 |
| GO:0043207 | response to external biotic stimulus | 2.9131788768940373e-6 |
| GO:0009607 | response to biotic stimulus | 3.3738092975918037e-6 |

**Supplementary Table 5.** Enrichment analysis of the set of genes identified as dynamic by scLANE but not by Lamian on the B-cell dataset [10]. The top-20 most-significant pathways by adjusted *p*-value are displayed.

#### 3 Supplementary Figures

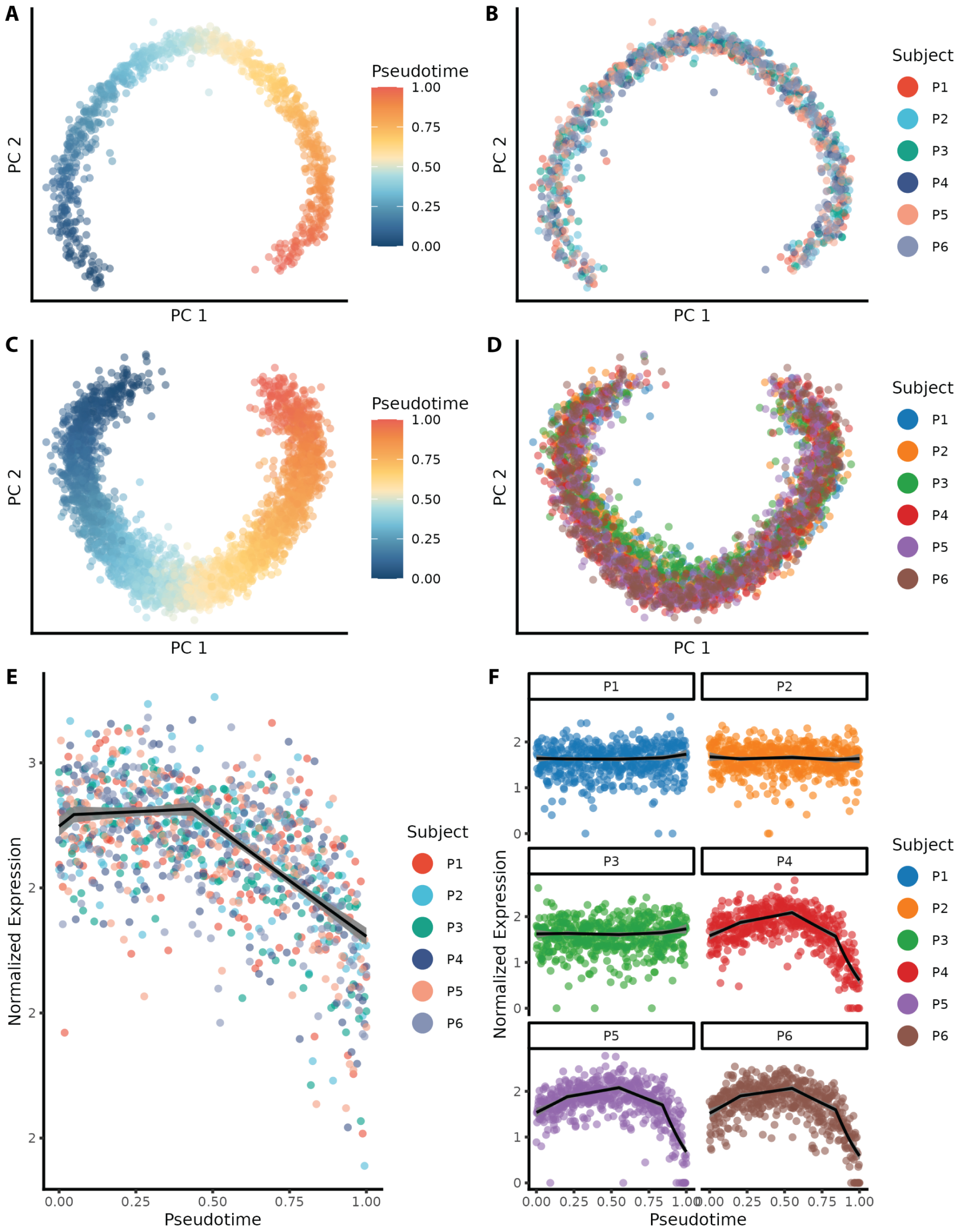


**Supplementary Figure 1. Simulation of realistic trajectory scRNA-seq data by Scaffold.** **(A)** PCA embedding of a representative multi-subject dataset upon which scLANE was run in GEE mode. Points correspond to individual cells, and are colored by ground-truth pseudotime. **(B)** PCA embedding of the cells from (A) with cells colored by subject ID. **(C)** PCA embedding of a representative multi-group, multi-subject dataset upon which scLANE was run in GLMM mode. Cells are colored by ground-truth pseudotime as in (A). **(D)** PCA embedding of the cells from (C) colored by subject ID. **(E)** Scatterplot of normalized expression of a truly-dynamic gene from the dataset shown in (A) with the GEE fit from scLANE overlaid in black. **(F)** Scatterplot of normalized expression of a truly-dynamic gene from the dataset shown in (C) with the per-subject GLMM fits from scLANE overlaid in black.

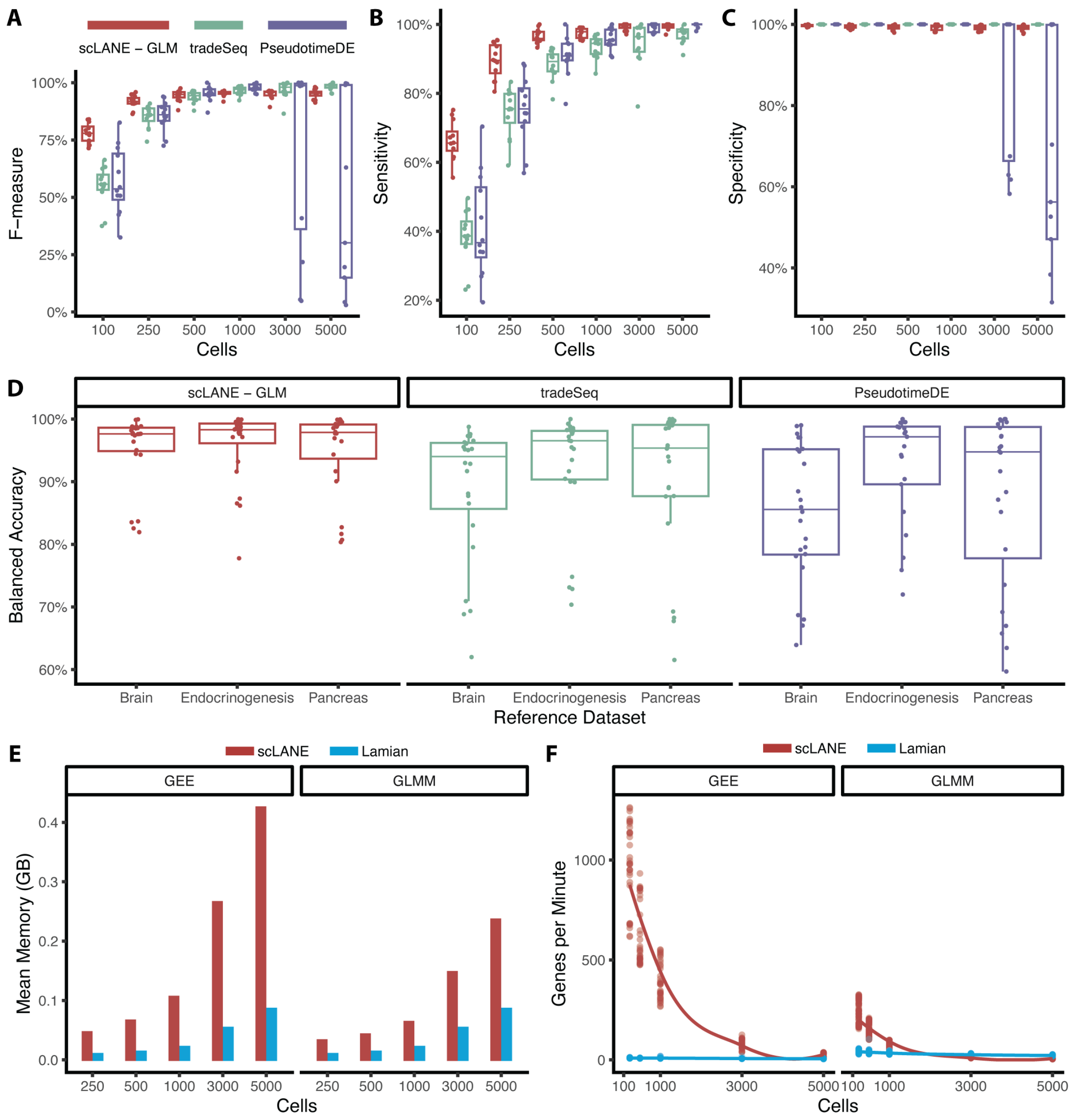


**Supplementary Figure 2. scLANE exhibits superior performance on simulated datasets.** **(A)** Boxplots of F-measure scores by number of cells for scLANE (in GLM mode), tradeSeq, and PseudotimeDE. **(B)** Similar to (A) for sensitivity scores. **(C)** Similar to (A) for specificity scores. **(D)** Boxplots of balanced accuracy by reference dataset for scLANE-GLM, tradeSeq, and PseudotimeDE. **(E)** Barplots of memory usage in gigabytes for the GEE and GLMM modes of scLANE and Lamian. **(F)** Similar to (E) for processing speed as measured by genes per minute.

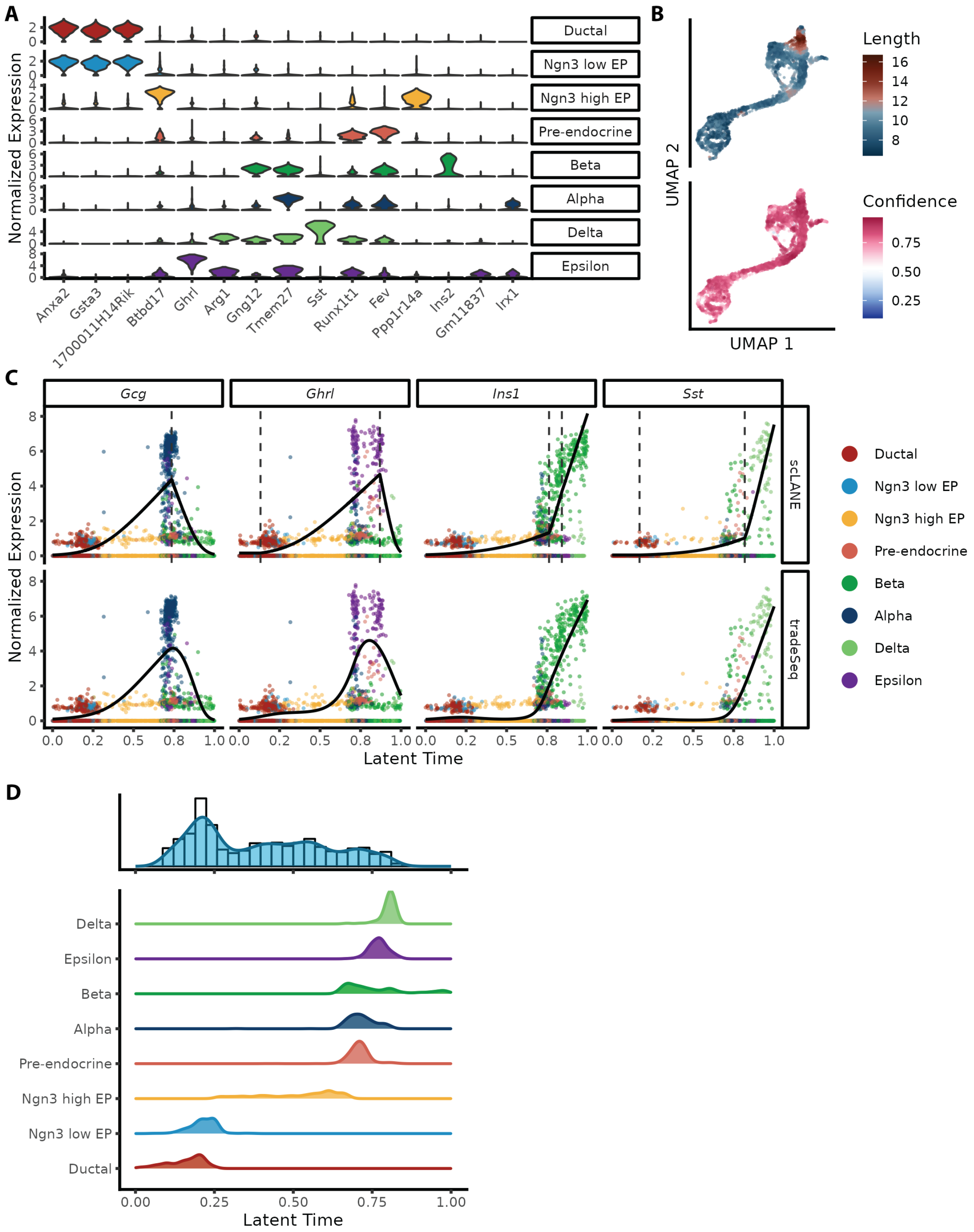


**Supplementary Figure 3. Characterizing gene dynamics in the pancreatic endocrinogenesis dataset. (A)** Violin plots of normalized expression for the top 2 marker genes by Wilcox test for each celltype in the dataset. **(B)** UMAP embedding colored by velocity length (top) and confidence (bottom). Velocity vector length is a proxy measurement for differentiation speed. **(C)** Fitted dynamics from scLANE and tradeSeq for the four peptide-encoding genes expressed by each mature endocrine celltype. Knots identified by scLANE are represented by dashed lines. **(D)** Empirical distribution and histogram of knots identified by scLANE, with ridgeline density plots of the distribution of latent time per celltype displayed below.

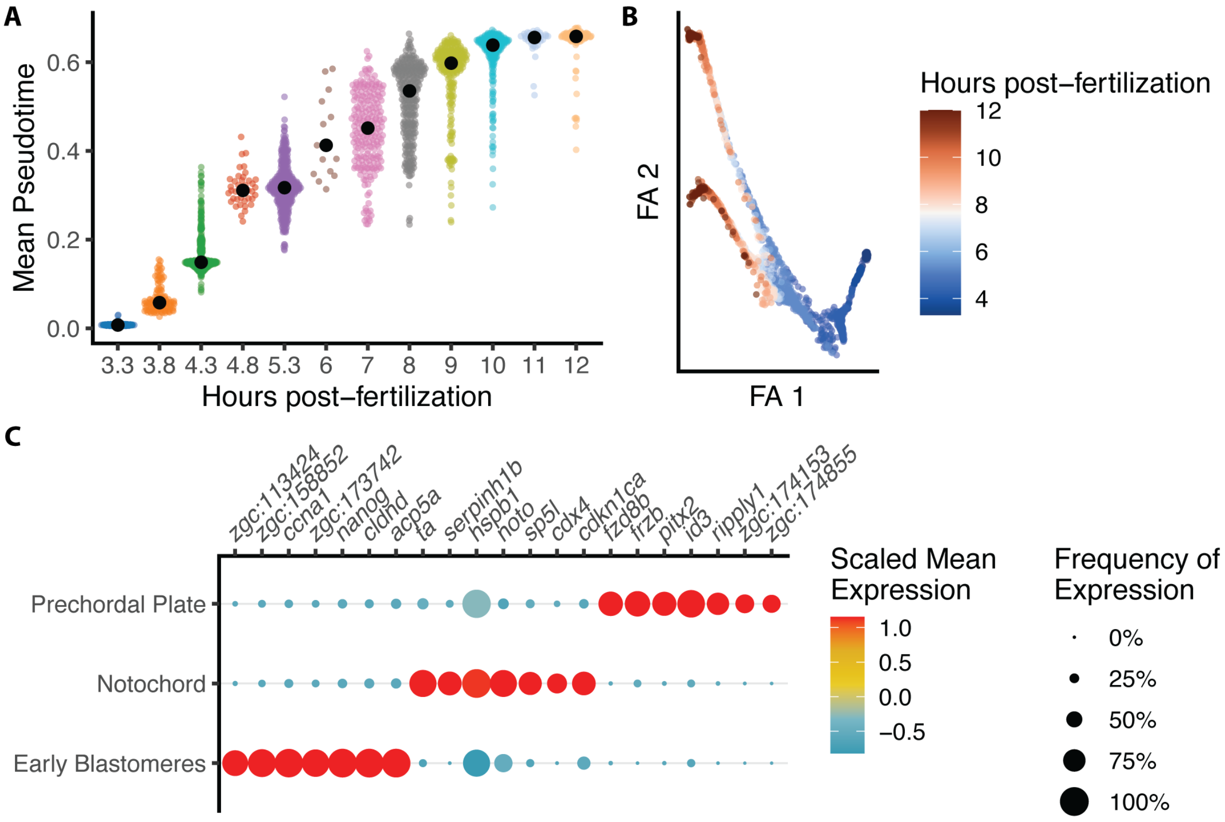


**Supplementary Figure 4. Examining differences between lineages in the time-resolved zebrafish dataset.**
*(A)* Beeswarm plots of mean-aggregated pseudotime by experimental timepoint, showing that Slingshot pseudotime has accurately recapitulated real experimental time. **(B)** Force-directed graph embedding colored by experimental time. **(C)** Dotplot displaying differences in expression for the top seven marker genes by celltype via Wilcox test. Color indicates scaled mean expression across celltypes, while radius denotes frequency of nonzero gene expression.

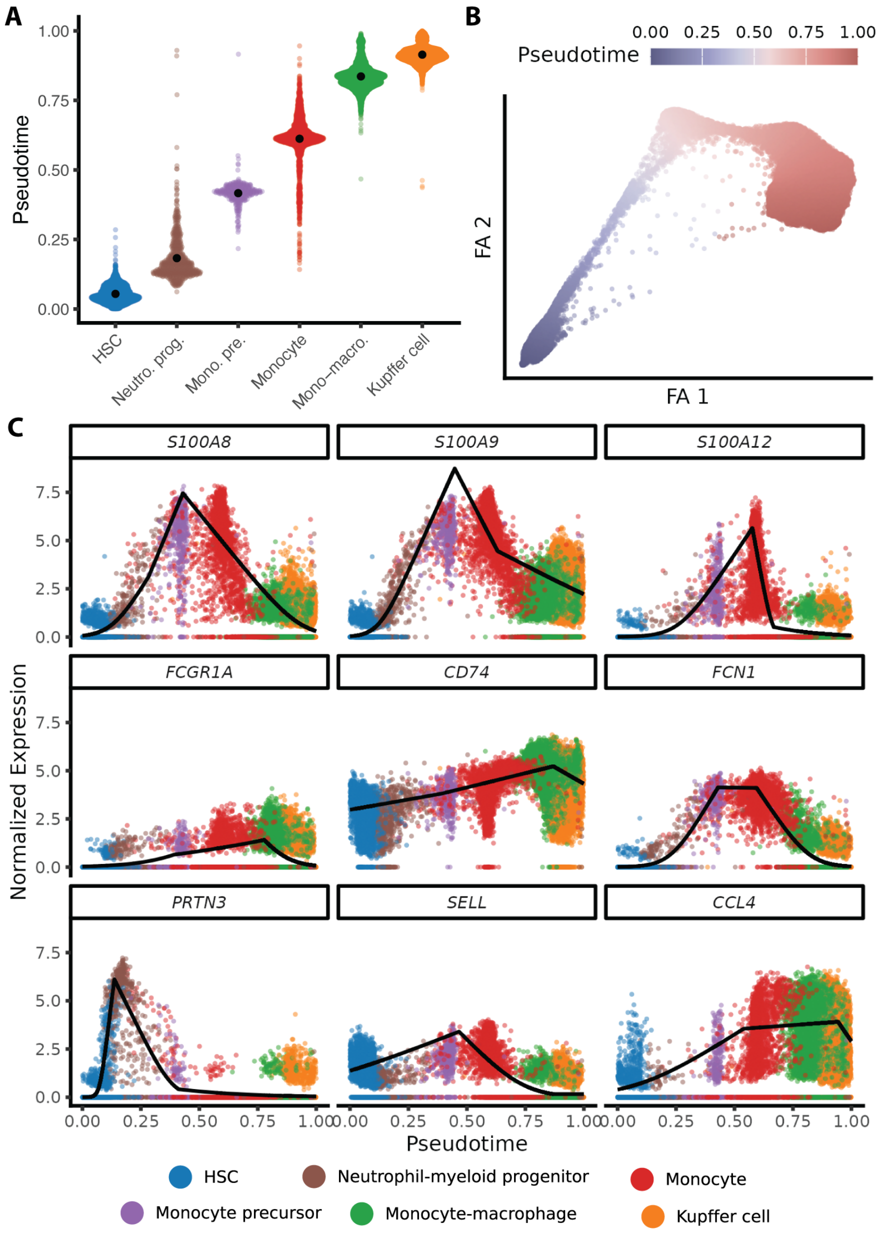


**Supplementary Figure 5. scLANE identifies biologically relevant gene programs regulating hematopoesis. (A)** Beeswarm plots of the distribution of pseudotime for each celltype. **(B)** Force-directed graph embedding colored by Slingshot-derived pseudotime. **(C)** Scatterplots of normalized gene expression over pseudotime for nine genes of interest from the original paper. The dynamics from scLANE are overlaid in black.

#### 4 Supplementary Source Code

```r
library(scLANE)
library(Seurat)
data_url <- url("https://zenodo.org/records/10012311/files/seu_panc.Rds")
seu_panc <- readRDS(data_url)
cell_offset <- createCellOffset(seu_panc)
lt_df <- data.frame(LT = seu_panc$latent_time)
candidate_genes <- chooseCandidateGenes(seu_panc,
                                        group.by.subject = FALSE,
                                        n.desired.genes = 3000L)
scLANE_models <- testDynamic(seu_panc,
                             pt = lt_df,
                             genes = candidate_genes,
                             size.factor.offset = cell_offset,
                             n.cores = 8L)
scLANE_de_res <- getResultsDE(scLANE_models)
```

**Supplementary Source Code 1. Performing scLANE testing in R.** Assuming the necessary packages have already been installed, this code allows the user to download the processed pancreatic endocrinogenesis dataset [7, 11], create the necessary inputs (a sequencing depth offset, dataframe containing the latent time ordering, and a list of genes to model), run the scLANE model, and summarize the results in a tidy table.

```r
library(dplyr)
dyn_genes <- filter(scLANE_de_res, Gene_Dynamic_Overall == 1) %>%
             pull(Gene)
scLANE_knots <- getKnotDist(scLANE_models, dyn.genes = dyn_genes)
dynamics_matrix <- smoothedCountsMatrix(scLANE_models,
                                        size.factor.offset = cell_offset,
                                        pt = lt_df,
                                        genes = dyn_genes,
                                        log1p.norm = TRUE)

library(bluster) #BiocManager::install("bluster")
library(coop) #install.packages("coop")
gene_embedding <- embedGenes(dynamics_matrix$Lineage_A)
library(UCell) #BiocManager::install("UCell")
seu_panc <- geneProgramScoring(seu_panc,
                               genes = gene_embedding$gene,
                               gene.clusters = gene_embedding$leiden)
library(gprofiler2) #install.packages("gprofiler2")
dyn_gene_enrich <- enrichDynamicGenes(scLANE_de_table, species = "mmusculus")
```

**Supplementary Source Code 2. Downstream analysis of scLANE results.** Again assuming that the necessary packages have been installed, and that the code fitting the scLANE models has been run, this code details several downstream analysis functionalities implemented in our package. After identifying the set of dynamic genes, a table of unique knot values is retrieved. Next, a matrix of smoothed gene dynamics is generated, after which a gene-level embedding of said dynamics is estimated using UMAP [3, 4] and clustered using the Leiden algorithm [12]. The resulting clusters of dynamic genes are termed dynamic gene programs, and cell-level scoring for each program is performed using UCell [13]. Finally, pathway enrichment analysis is performed for the set of dynamic genes.

```r
library(dplyr)
library(scLANE)
library(Seurat)
library(tradeSeq)
data_url <- url("https://zenodo.org/records/10012311/files/seu_bcell.Rds")
seu_bcell <- readRDS(data_url)
candidate_genes <- chooseCandidateGenes(seu_bcell,
                                        id.vec = seu_bcell$fetal.ids,
                                        n.desired.genes = 3000L)
RNA_counts <- as.matrix(seu_bcell@assays$RNA$counts)[candidate_genes, ]
cell_offset <- createCellOffset(seu_bcell)
pt_df <- data.frame(DPT = seu_bcell$dpt_pseudotime)
bioc_par <- BiocParallel::MulticoreParam(workers = 24L, RNGseed = 312)
k_eval <- evaluateK(RNA_counts,
                    pseudotime = pt_df,
                    cellWeights = matrix(rep(1, nrow(pt_df)), ncol = 1),
                    offset = log(1 / cell_offset),
                    k = 3:10,
                    plot = FALSE,
                    nGenes = 500,
                    verbose = FALSE,
                    parallel = TRUE,
                    BPPARAM = bioc_par)
best_k <- c(3:10)[which.min(abs(colMeans(k_eval - rowMeans(k_eval))))]
ts_models <- fitGAM(RNA_counts,
                    pseudotime = pt_df,
                    cellWeights = matrix(rep(1, nrow(pt_df)), ncol = 1),
                    offset = log(1 / cell_offset),
                    nknots = best_k,
                    sce = FALSE,
                    parallel = TRUE,
                    verbose = FALSE,
                    BPPARAM = bioc_par)
BiocParallel::bpstop(bioc_par)
ts_de_table <- associationTest(ts_models, global = TRUE) %>%
               arrange(desc(waldStat)) %>%
               mutate(gene = rownames(.),
                      pvalue_adj = p.adjust(pvalue, method = "fdr"),
                      gene_dynamic_overall = if_else(pvalue_adj < 0.01,
                                                     1,
                                                     0)) %>%
               relocate(gene)
```

**Supplementary Source Code 3. Performing tradeSeq testing in R.** Assuming that the necessary packages have been installed, this code allows the user to download the processed B-cell maturation dataset [10], then run the tradeSeq method [14]. First, a counts-per-10k sequencing depth offset is created using the `createCellOffset()` function from scLANE. Next, a dataframe containing diffusion pseudotime is provided along with expression counts and the offset to the `evaluateK()` function, then the “best” number of knots $k$ is selected by identifying the value with the lowest mean absolute deviation from the mean AIC. Finally, the models are fitted, and then summarized using a global Wald test for association with pseudotime.

```
library(dplyr)
library(scLANE)
library(Seurat)
library(Lamian)
data_url <- url("https://zenodo.org/records/10012311/files/seu_bcell.Rds")
seu_bcell <- readRDS(data_url)
candidate_genes <- chooseCandidateGenes(seu_bcell,
                                        id.vec = seu_bcell$fetal.ids,
                                        n.desired.genes = 3000L)
cell_anno <- select(seu_bcell@meta.data,
                    Cell = cell,
                    Sample = fetal.ids) %>%
             mutate(across(everything(), as.character)) %>%
             magrittr::set_rownames(NULL)
cell_pt <- as.integer(rank(seu_bcell$dpt_pseudotime))
names(cell_pt) <- cell_anno$Cell
subject_intercept <- rep(1, length(unique(seu_bcell$fetal.ids)))
samp_design <- data.frame(intercept = subject_intercept) %>%
               magrittr::set_rownames(unique(seu_bcell$fetal.ids)) %>%
               as.matrix()
RNA_counts <- as.matrix(seu_bcell@assays$RNA$data)[candidate_genes, ]
lamian_models <- lamian_test(RNA_counts,
                             cellanno = cell_anno,
                             pseudotime = cell_pt,
                             design = samp_design,
                             test.type = "time",
                             test.method = "permutation",
                             permuiter = 100,
                             verbose.output = FALSE,
                             ncores = 24L)
```

**Supplementary Source Code 4. Performing Lamian testing in R.** Assuming that the necessary packages have been installed, this code allows the user to download the processed B-cell maturation dataset [10], then run the Lamian method [15]. First, a dataframe containing per-cell annotations is generated along with a vector of integer-valued diffusion pseudotime estimates and a matrix specifying the experimental design. Next, the `lamian_test()` function is used to perform permutation-based testing of each gene's normalized expression with pseudotime.

[3] Leland McInnes, John Healy, and James Melville. "UMAP: Uniform Manifold Approximation and Projection for Dimension Reduction". In: (Sept. 2020). Publisher: arXiv Version Number: 3.

[4] James Melville. *uwot: The Uniform Manifold Approximation and Projection (UMAP) Method for Dimensionality Reduction.* 2022.
